# Spatial frequency-based correction of the spherical aberration in living brain imaging

**DOI:** 10.1101/2022.12.05.519048

**Authors:** Aoi Gohma, Naoya Adachi, Daiki Horiba, Yasuo Yonemaru, Daisuke Nishiwaki, Eiji Yokoi, Kaori Higuchi, Yoshihiro Ue, Atsushi Miyawaki, Hiromu Monai

**Affiliations:** Graduate School of Humanities and Sciences, Ochanomizu University, Ohtsuka, Bunkyo-ku, Tokyo, 112-8610, Japan; RIKEN Center for Brain Science-Evident Open Collaboration Center (BOCC), 2-1, Hirosawa, Wako-shi, Saitama, 351-0106, Japan

**Keywords:** Objective lens, Correction collar, Refractive index

## Abstract

We previously developed a fully automated spherical aberration compensation microscope system, Deep-C, to obtain spherical aberration-free images, but the contrast-based algorithm (Peak-C) may limit applications for low signal-to-noise ratio images. Herein we propose a new spatial frequency-based algorithm called Peak-F and compared its performance to Peak-C. Unlike Peak-C, Peak-F is robust to any noise level since it is independent of the dynamic range of the images, and it does not suffer from image saturation. Finally, Peak-F was implemented in a two-photon microscope to observe living aged and young mouse brains. Consequently, the average refractive index of brain tissue was higher in old mice than in young mice. The Peak-F algorithm determines high-resolution microscopic images stably and robustly.

## Introduction

Optical errors often hamper high-resolution optical imaging of biological samples. Such errors are due to biochemical components such as water, proteins, and lipids as well as physical properties such as anisotropy and the refractive index (RI) of the biological tissue. Additionally, induced tissue scattering limits image acquisition in deeper regions or aged tissue. Among the optical errors, spherical aberrations are due to mismatched RIs between the immersion medium (e.g., water) and the sample, causing photons to converge into two different focal planes[1–5]. A correction collar, which is usually attached to the objective lens with a high numerical aperture, is a revolving optical mechanism to reduce the mismatch between RIs[6,7]. It adjusts the distance of the inner lens to correct spherical aberrations. In the case of a histological sample, for example, the correction collar is adjusted to “θ = 0 degrees” (indicating 0.17 μm corresponds to the thickness of the cover glass). However, changing the correction collar is difficult for thick samples because there is not an objective measurement of the necessary rotation to eliminate spherical aberrations. To obtain spherical aberration-free images, we developed a fully automated spherical aberration compensation microscope system called Deep-C[8]. This system has a motorized correction collar, and the contrast of the entire image quantifies spherical aberrations at a given rotation angle.

A contrast-based calculation is often used to determine the sharpness of an image to maximize its contrast value since a clear image has a high contrast value, whereas a blurred one has a low contrast value. For example, out-of-focus (no focus), an optical phenomenon, occurs when multiple subjects are at different distances from the lens and the light passing through the lens forms images at various locations. It is impossible to have all images in focus on the same focal plane simultaneously. Hence, unfocused light sources produce unclear images for the observer. Auto-focusing involves adjusting the distance between the lens and the specimen to focus the light automatically[9]. To determine the optimal focal length, the Brenner gradient method is widely used to maximize contrast[10]. In this method, the total contrast is equal to the square of the difference in intensity between neighboring pixels, which is given as

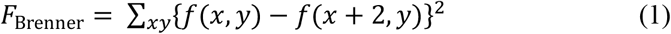

Although different physical phenomena cause spherical aberrations and out-of-focus, they are similar since the resulting light does not focus on a single point. Therefore, spherical aberrations may be corrected using a method to resolve blurriness in an out-of-focus image. Unfortunately, certain frequency bands cannot be evaluated by simply adapting the Brenner gradient to microscope images due to the mismatch between the contrast evaluation region and the spatial frequency of the image. We solved this problem by calculating the contrast value as the sum of the squared differences among 1, 2, 3, 5, and 10 neighbors. Our approach, called Peak-C, accommodates a wide range of spatial frequencies.

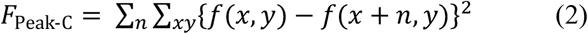

Our previous system consisted of a motorized collection collar and Peak-C algorithm, which calculated the optimal rotation[8]. We demonstrated fine and quantitative two-photon images, for example, of dendritic spines with this system. However, testing of the Peak-C algorithm indicated that biological scattering may result in many false positives. Thus, applying Peak-C to low signal-to-noise ratio images such as the deep brain areas and aged brain tissues is challenging.

Our research aims to understand Peak-C’s susceptibility to noise and propose an alternative approach. Herein we propose a spatial frequency-based system, Peak-F, to find the optimum correction collar position from images with noise. Peak-F is a spatial-frequency bandpass filter based on a fast Fourier transform that extracts low-frequency boundaries from the images. This approach is based on the observation that noise appears in the high-frequency region. Comparing Peak-C and Peak-F reveals that Peak-F possesses a noise-resilient nature and is stable to other disturbances such as saturation. Furthermore, we incorporated Peak-F into a two-photon microscope, allowing the deep area of an aged mouse brain to be imaged.

## Materials and Methods

### Surgical procedures

Benchmark images were obtained from a previous study[8]. Male and female Thy1-YFP-H(YFP-H) transgenic mice[11] were used. Young mice were 9-, 11-, 25-, and 25-weeks of age, while old mice were 52-, 81-, 83-, and 91-weeks of age. Mice were housed under a 12-h:12-h light:dark cycles and raised in groups up to five. Mice were anesthetized with isoflurane (2%), and their body temperature was maintained at 37 °C with a heating pad (BWT-100 A, Bio Research Center or TR-200, Fine Science Tools) during surgery and recording. After skull exposure, a metal frame was attached to the skull using dental acrylic (Fuji LUTE BC, GC, Tokyo, Japan; Super Bond C&B, Sunmedical, Shiga, Japan). For two-photon imaging, a craniotomy (2.7-mm diameter) was made above the visual cortex (AP −2.0 mm, ML +2.5 mm). Then the dura mater was surgically removed. Next, the craniotomy was gently sealed with a thin glass coverslip (2.7 × 2.7 mm, thickness: 0.12–0.17 mm, Matsunami Glass, Osaka, Japan). Finally, the cranial window was secured with dental cement (Fuji LUTE BC, GC, Tokyo, Japan; Super Bond C&B, Sunmedical, Shiga, Japan).

### Two-photon excitation imaging

Multi-photon laser scanning microscope (FVMPE-RS, Evident) equipped with an InSight laser system (Spectra-Physics, 960-nm wavelength) and an Olympus objective (FV30-AC25W, NA: 1.05, working distance: 2 mm, immersion medium: water) was used. In Figs. 1–4, the image size is 1024 × 1024 pixels (16-bit resolution), while the image size in Fig. 5 is 1024 × 1024 pixels (8-bit resolution).

**Fig. 1.**
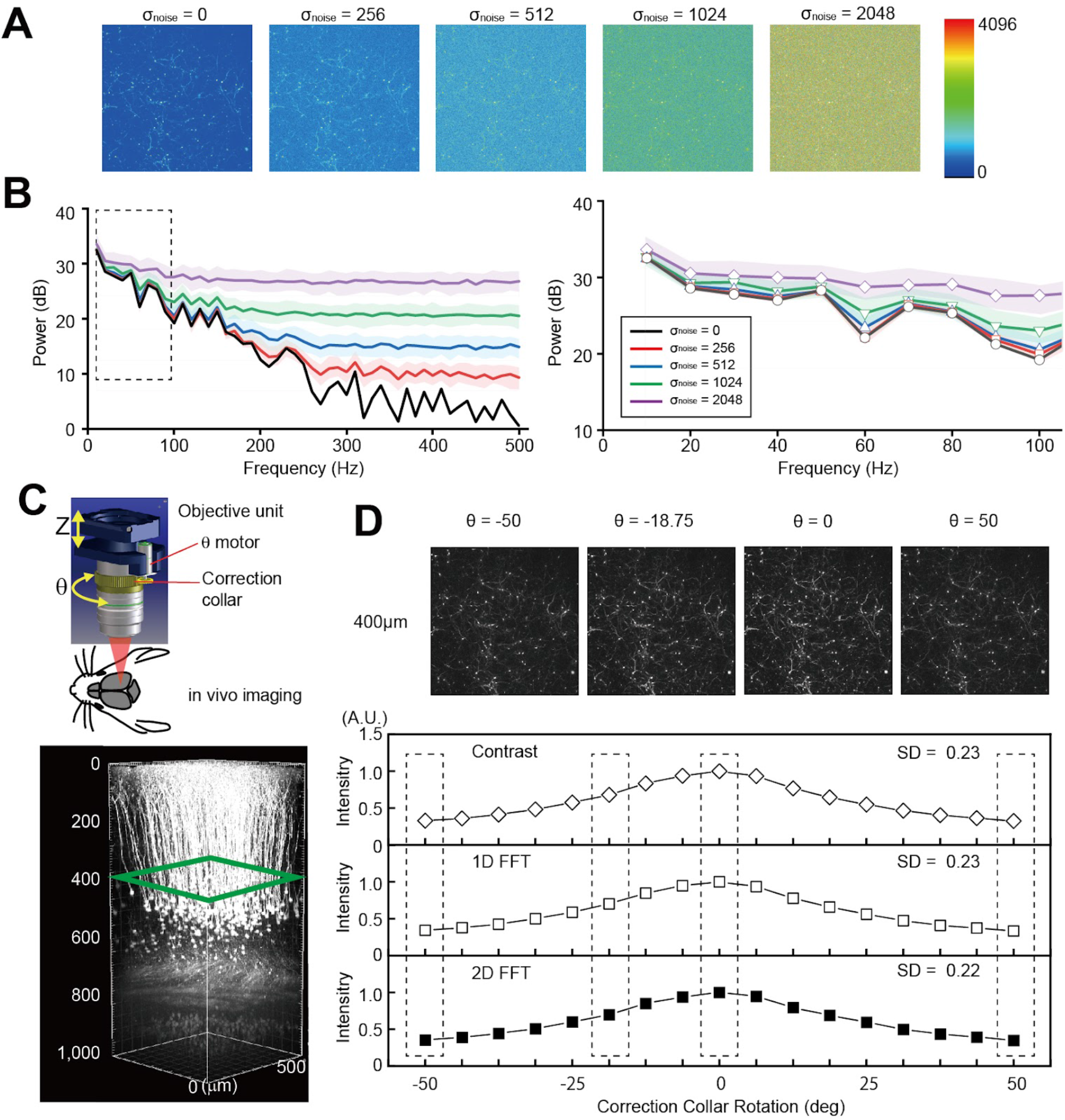
Determination of the optimal correction collar position based on the image spatial frequency A. Benchmark image and its image with noise added (σ = 256, 512, 1024, and 2048) B. Power spectrum for σ = 256(red curve), 512(blue curve), 1024(green curve), and 2048(purple curve). The power spectrum for the frequency range 0 - 40 Hz is shown in the dashed square (right). The vertical axis is Power, the horizontal axis is frequency, and the colored area is the standard error. C. Objective lens position and observed area. D. Variation of contrast intensity and power intensity with the angle of the correction collar. The vertical axis is the contrast or power intensity, and the horizontal axis is the rotation angle of the correction collar. The traditional method is white diamonds, FFT1 is white squares, and FFT2 is black squares.

**Fig.2.**
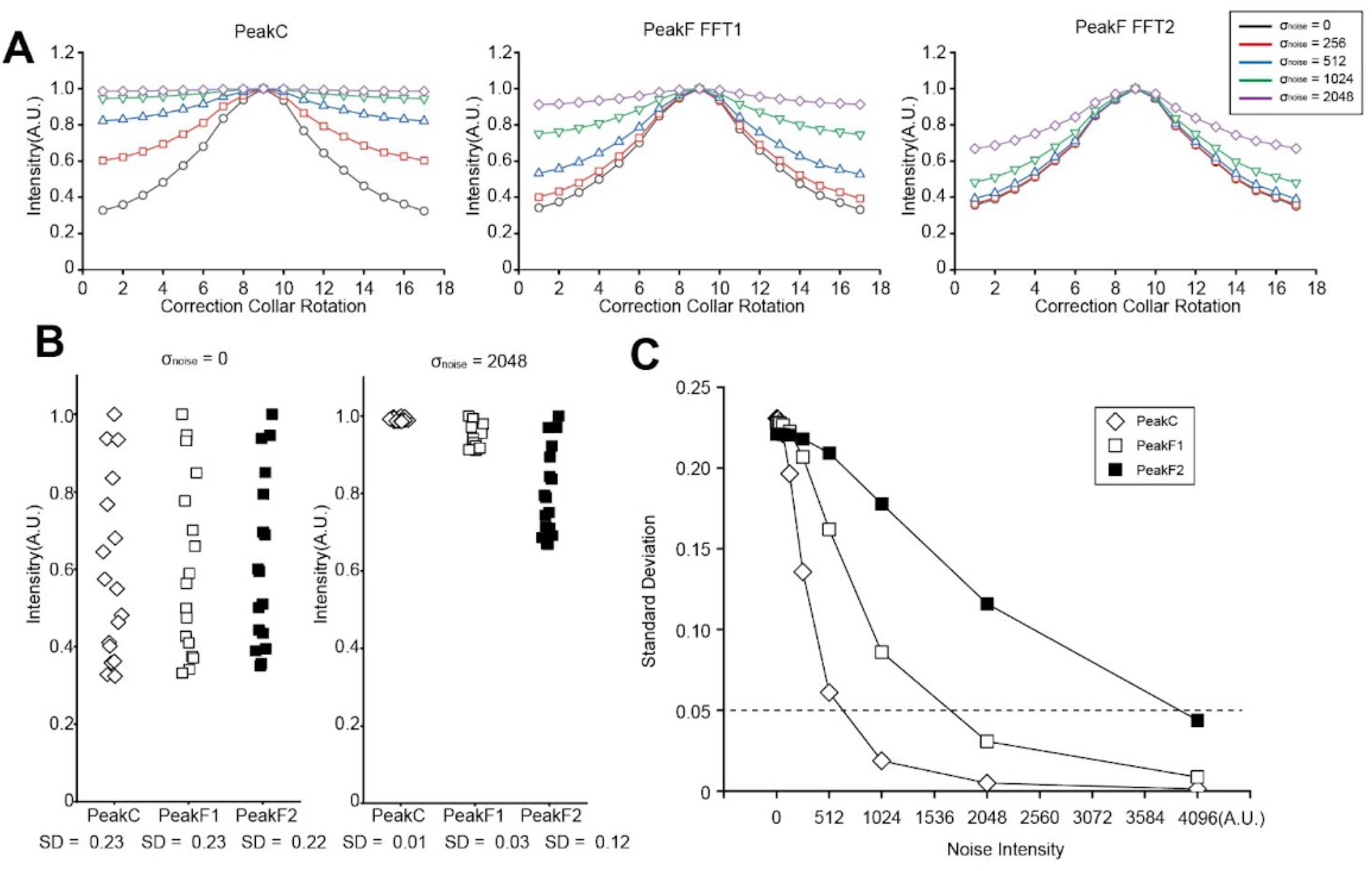
Peak-F algorithm is robust to noise A. Intensities at each of the 17 correction collar positions calculated by the Peak-C, Peak-F FFT1, and Peak-F FFT2 algorithms for images with noise added at σ = 256 (red), 512 (blue), 1024 (green), and 2048 (purple). Intensities are normalized to the maximum value. B. The ability to detect peaks with no noise (σ_noise = 0) and strong noise (σ_noise = 2048) was evaluated by the variation (standard deviation) of values at 16 locations. Peak-C is represented by a white diamond, Peak-F FFT2 by a white square, and Peak-F FFT2 by a black square. C. Trends in the value of standard deviation for each noise intensity. Peak-C is represented by a white diamond, Peak-F FFT2 by a white square, and Peak-F FFT2 by a black square.

**Fig.3.**
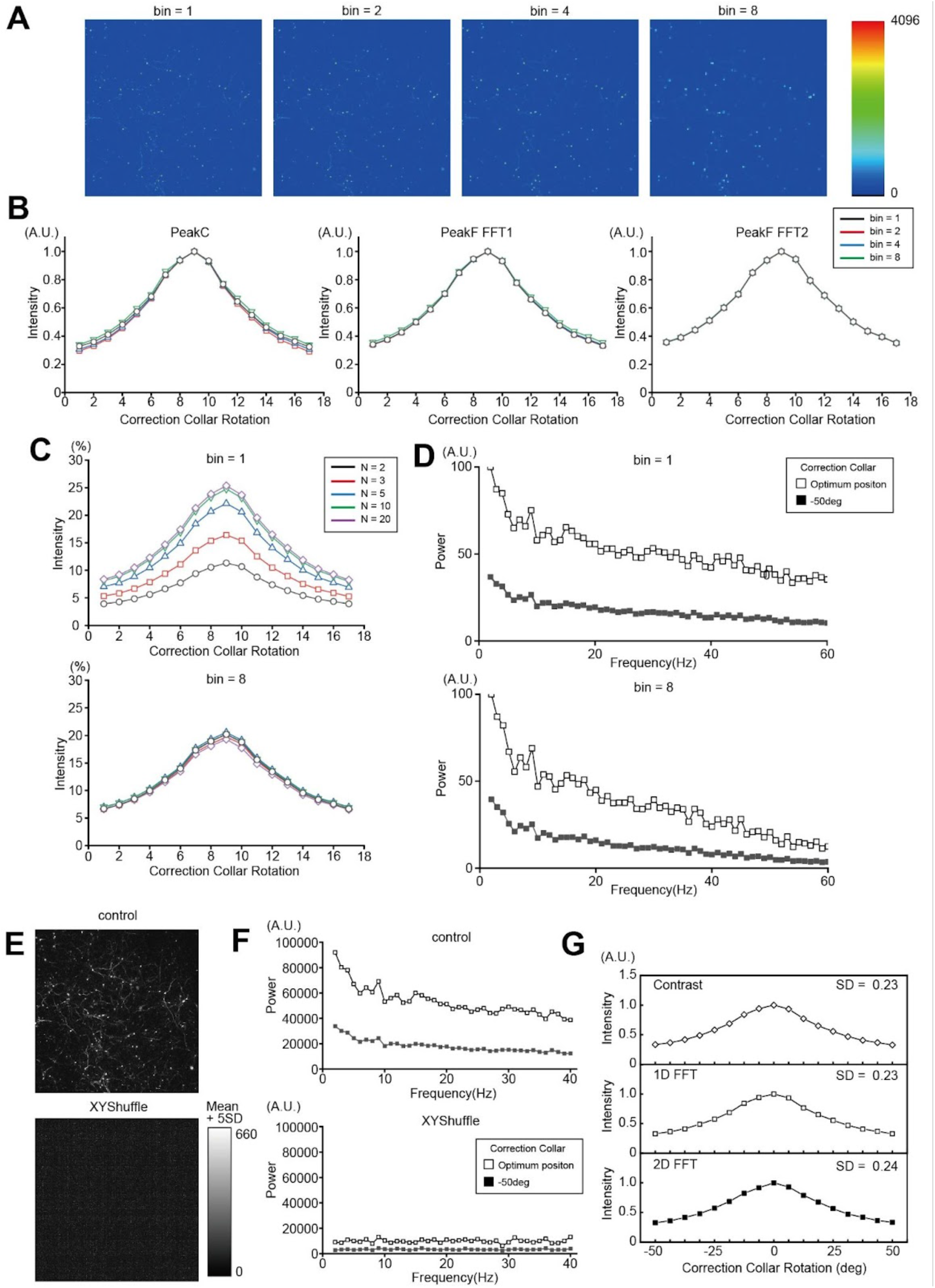
Comparison of Peak-C’s and Peak-F’s performance on noise-added images and binning images A. Binning was performed on the benchmark image. 256 × 256 (bin = 2), 128 x 128 (bin = 4), 64 × 64 pixel (bin = 8) images were created. B. Intensities for each of the 17 correction collar positions calculated with the Peak-C, Peak-F FFT1, and Peak-F FFT2 algorithms for the image A. bin = 1 is black, bin = 2 is red, bin = 4 is blue, and bin= 8 is green. The vertical axis is the contrast or power intensity, the horizontal axis is the rotation angle of the correction collar, and the intensity is normalized by the maximum value. C. The contribution of the Peak-C algorithm at each N when bin = 1 and bin = 8. N = 2 is black, N = 3 is red, N = 5 is blue, N = 10 is green, and N =20 is purple. The vertical axis is the intensity of contrast or power, the horizontal axis is the correction collar rotation angle, and the intensity is normalized by the maximum value. D. Power spectrum of the image at bin = 1 and bin = 8. The vertical axis is power, the horizontal axis is frequency, and the vertical axis is normalized by the maximum value. Calculations were performed with FFT1. The graph of the optimal correction ring position is represented by a white square and the graph of the improper correction ring position by a black square. E. XY shuffled image. For ease of viewing, the threshold is set to the average of the actual brightness + 5 SD, and all pixel brightness above the threshold are set to 660. F. Power spectrum of the reference image and the XY shuffled image. The vertical axis is power and the horizontal axis is frequency, with the vertical axis normalized by the maximum value. Calculations were performed with FFT1. The graph of optimal correction ring positions is represented by white squares and the graph of inappropriate correction ring positions by black squares. G. The intensity of each of the 17 correction collar positions was calculated with the Peak-C (white diamond), Peak-F FFT1 (white square), and Peak-F FFT2 (black square) algorithms on the A image. The vertical axis is the contrast or power intensity, the horizontal axis is the rotation angle of the correction collar, and the intensity is normalized by the maximum value.

**Fig.4.**
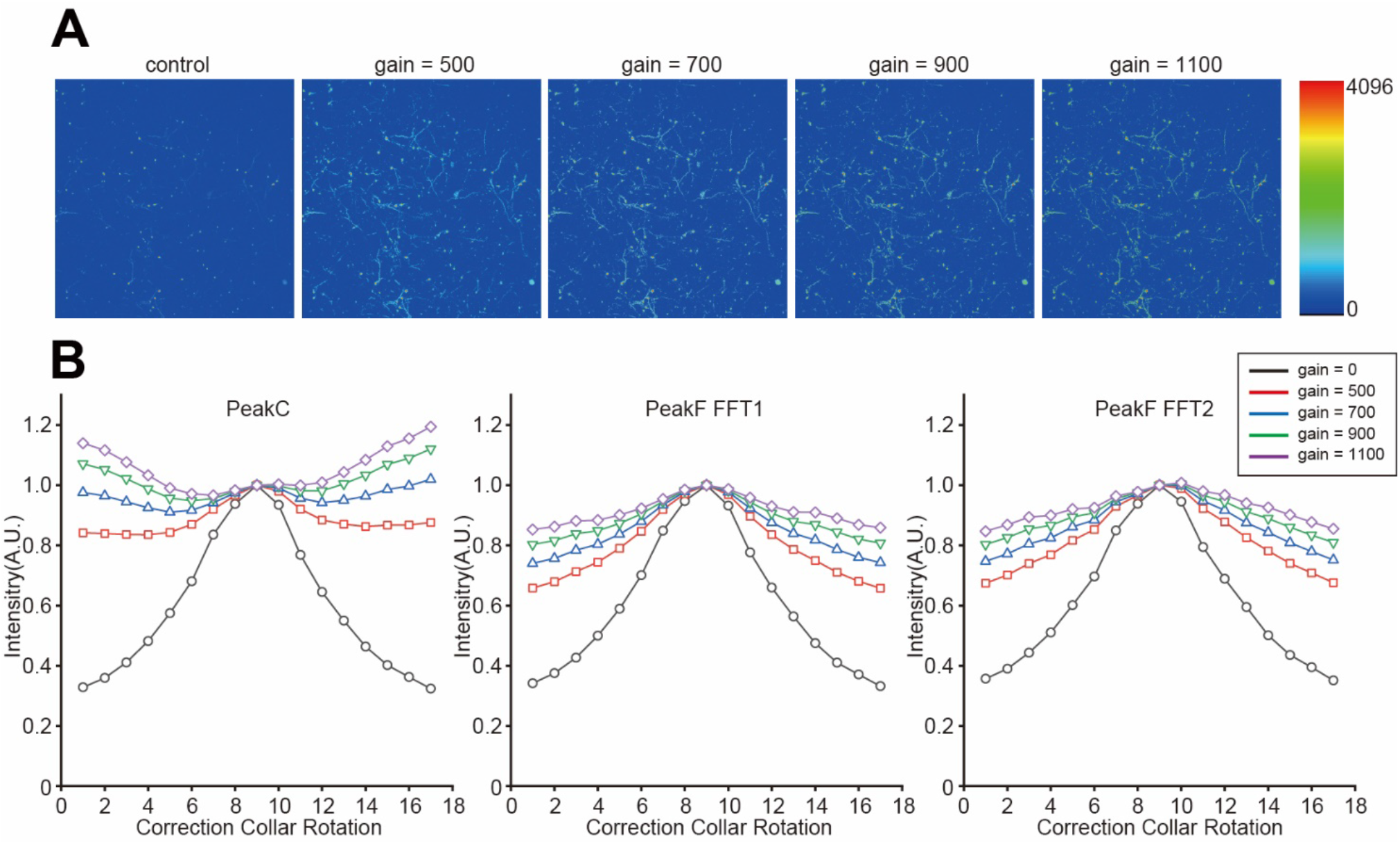
Comparison of Peak-C’s and Peak-F’s performance on saturation-added images A. Benchmark image and its image with four levels of saturation (gain=500, 700, 900, and 1100). B. Intensity of each of the 17 correction ring positions calculated by the Peak-C, Peak-F FFT1, and Peak-F FFT2 algorithms when saturation was applied to the benchmark image. The vertical axis is the contrast or power intensity, the horizontal axis is the rotation angle of the correction collar, and the intensity is normalized by the maximum value.

**Fig.5.**
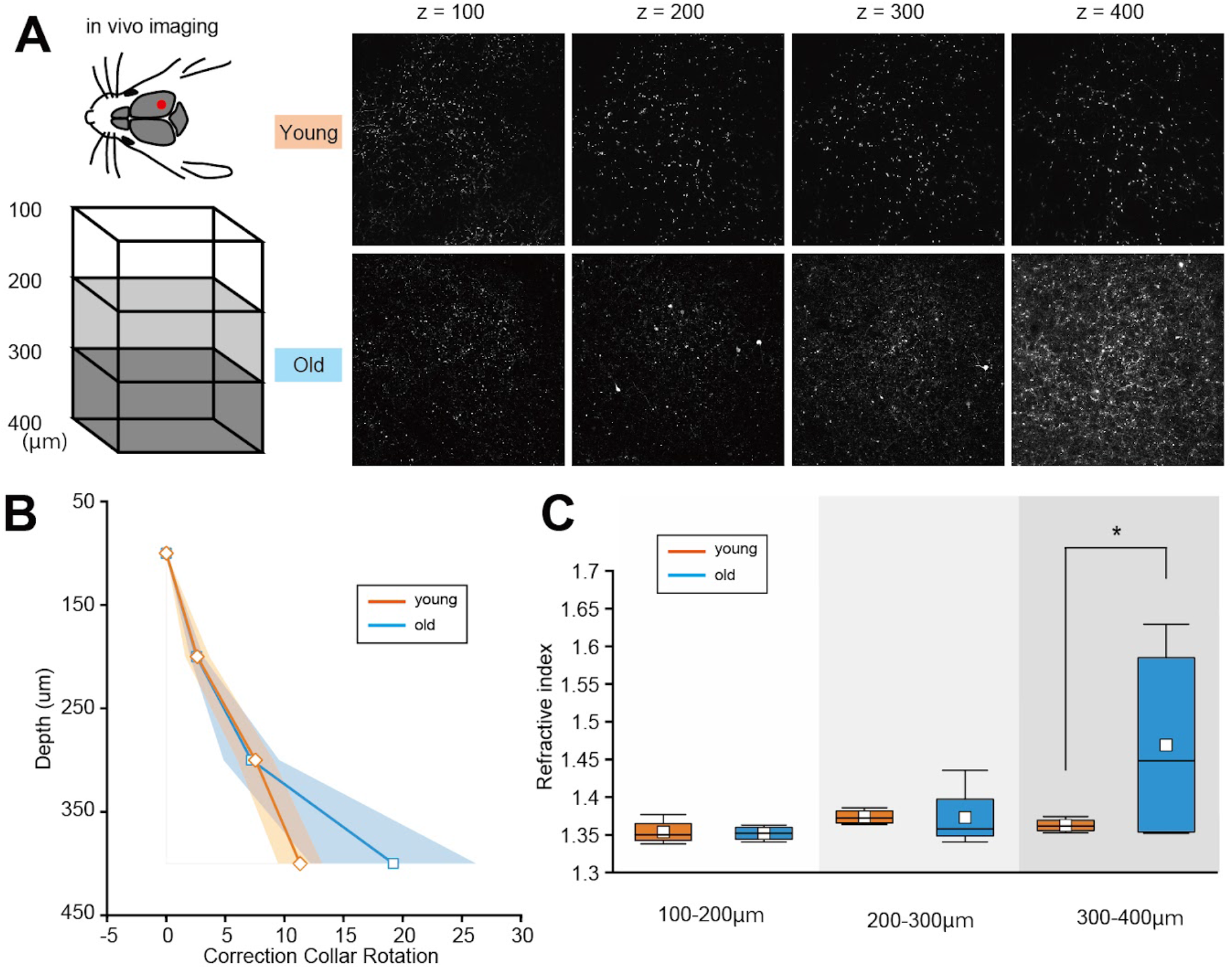
Peak-F reveals optical characteristics of the brain tissue from noisy images A. Images of 100, 200, 300, and 400 μm below the surface of the brain of young and old mice. B. Mean (solid line) and standard deviation (colored area) of the optimal angle of the correction collar for young and old mice. C. The tissue refractive index was calculated from the optimal angle of the correction collar for young and old mice. 300um-400um showed a significant difference between young and old mice. Tests were performed with Two-way ANOVA. *p < 0.05.

### Data analysis

The acquired Fluoroview .oir file was converted into a 1024 × 1024 pixel, 8-bit grayscale bitmap image. Then the calculations were performed using MATLAB.

#### Peak-C

The Peak-C algorithm was calculated using the formula from the previous study[8].

#### Peak-F 1D FFT

Fast Fourier transforms an image one column at a time from the top as a matrix. Then it determines the optimal angle of the correction collar by comparing the total intensity of the extracted power spectrum from the frequencies ranging from 2 to 40 Hz for each correction collar position.

#### Peak-F 2D FFT

2-D fast Fourier transforms the entire image. Then it determines the optimal angle of the correction collar by comparing the total intensity of the extracted power spectrum from frequencies ranging from 2 to 40 Hz for each correction collar position.

#### Adding white noise

A 1024 × 1024 random matrix was generated using the Matlab function rand. Then multiplying the random matrix by σ created noise images for several intensities. The noise was reproduced by adding the benchmark image and the noise image created. σ was set to 256, 512, 1024, or 2048. Additionally, σ was set to 0 or as a control.

#### Binning

Bicubic interpolation was performed on the benchmark image. The matrix size was reshaped to 1024 × 1024 for bin=1, 512 × 512 for bin=2, 128 × 128 for bin=4, and 64 × 64 for bin=8.

#### Random shuffle

The rows and columns of each pixel in the benchmark image were randomly placed using the function randperm.

#### Adding saturation

The mean value + the standard deviation (SD) of the luminance of the benchmark image was used as the threshold value. Then 500, 700, 900, or 1100 were added to the values of pixels above the threshold value. Afterwards, pixel values above 4095 were set to 4095.

### Implementation to two-photon microscope

Peak-F calculations for two-photon excitation imaging were performed by modifying the FV31S-SW software of the FVMPE-RS TruResolution (Evident) system. Specially, the contrast calculation processing part of Peak-C was replaced with Peak-F. The image frequency was calculated by replacing only the contrast calculation part of Peak-C with the image frequency calculation process of Peak-F. Image frequencies were calculated using the well-known Cooley-Tukey type Fast Fourier Transform method.

## Results

### Peak-F algorithm is robust to noise

Peak-F was designed in two ways. One spatial frequency-based algorithm named 1D FFT was the summation of the 1–40 Hz power spectrum of one line of the image. For example, when the image was taken by 1024-pixel × 1024-pixel, the power spectrum of the first line of the image was calculated and extracted by 2–40 Hz. This extraction process was repeated up to the 512th pixel and summed. The other is 2D FFT, which is a 2–40 Hz power spectrum of the whole image. The fast Fourier transform performed band-pass filtering (see Method).

Figure 1 explains why the chosen spectrum was 1–40 Hz. First, the noise is not negligible in the deep brain tissue, and we aimed to understand how noise affects the power spectrum of the spatial frequency calculated from typical in vivo brain images, which were acquired by a two-photon microscope (benchmark image, see Method). Then we added white noise with various intensities to the benchmark image (control, Fig. 1A).

As the noise intensity (σ_noise) increased, the effect of noise became non-negligible, especially in the high-frequency region, and the deviation from the power spectrum obtained from the reference image increased. In contrast, the low-frequency component below 40 Hz was less affected by noise (Fig. 1B). Consequently, a calculation frequency of 2–40 Hz was used in the Peak-F algorithm because it is less susceptible to noise and can accommodate a wide range of noise intensities. To verify whether the proposed Peak-F can determine the optimal correction collar position similar to Peak-C, we prepared images in which only the correction collar position was changed when acquiring the reference image, applied Peak-F, and tried to detect the image with maximum power at 1–40 Hz. In the images acquired at 400 μm from the brain surface of the cerebral cortex of Thy1-YFP H-line transgenic mice, 17 benchmark images were obtained by turning the motorized correction collar with a Deep-C objective from −50° to 50° at 6.25° intervals (Fig. 1C). Applying the Peak-C algorithm to the benchmark images indicated that θ = 0 is the optimal correction collar position because the maximum contrast is obtained when the rotation angle of the correction collar is 0 degrees (θ = 0) (Fig. 1D, upper). Similarly, Peak-F was applied to the waveform vectorized in the x-axis direction from the image. The results of Peak-F using 1D FFT and plotting the total power at 1–40 Hz showed that the maximum power was obtained when the rotation angle of the correction collar was 0 degrees (θ=0) (Fig.1D, middle). Applying 2D FFT gave the same results. Figure 1D (lower) plots the power values for the 2–40 Hz component.

The effect of noise is more pronounced after the image’s spatial frequency of 40 Hz (Fig. 1B). Therefore, we expect that Peak-F can stably determine the optimal correction collar position even in noise-added images. Next, we compared the abilities of Peak-C and Peak-F to detect the optimal correction ring position when the noise intensity was varied using the benchmark image created in Fig.1A with artificially added white noise (Fig. 2A). We determined the variation of contrast or power displacement of the images acquired at each correction ring position for noise intensity σ=0 and σ=2048. The difference in contrast required to detect the optimal correction ring position decreased as the noise intensity increased for each algorithm. However, the difference for Peak-F was more significant than Peak-C (Fig. 2B, SD = 0.01 vs. 0.03 vs. 0.12). An SD value less than 0.05 indicated that the system could not find the optimum position. Figure 2D shows the calculated SD for each noise intensity. The Peak-F algorithm using the two-dimensional FFT was particularly robust to noise, while the Peak-C algorithm was vulnerable.

### Peak-F algorithm finds the best images according to biological structures

Next, we investigated the difference in susceptibility to noise between the algorithms. Because we hypothesized the difference is due to the treatment of the spatial frequency, we prepared a binned image in which the spatial frequency was varied while preserving the image contrast (Fig. 3A). Peak-C found the optimal correction collar position even with binning only eight 8 pixels together (Fig. 3B). This suggests that Peak-C only considers image contrast.

As shown in Eq. (2), the Peak-C algorithm sums the contrast at each discrete spatial frequency (N = 2, 3, 5, 10, 20) to cover a wide range of spatial frequencies. This procedure increases the calculation cost. For Peak-C, nearly 75% of the contribution ratios had a large N value (N = 5, 10, or 20) (Fig. 3C, upper), indicating that the high-spatial-frequency domain (25–100 Hz) greatly contributes to determining the optimal correction collar position. However, since the effect of noise was non-negligible above 50 Hz (Fig. 1B), this may be why Peak-C is vulnerable to noise. In contrast, as the binning size increased, each N had similar contribution ratios (Fig. 3C, lower). This result suggests that in the case of binned images, calculations using N =2, which is known as Brenner’s equation (Eq. 1), are insufficient. Thus, the optimum spatial frequency should be chosen to reduce the calculation cost. However, it is usually difficult to assess the value of N. In contrast, Peak-F excluded the arbitrariness in selecting the optimal frequency in the calculation of any image because the frequency domain includes all spatial frequencies less than 40 Hz (Fig. 3D).

To consider the significance of the spatial-frequency distribution in biological structures, we generated a randomly XY-shuffled image in which the image had the same contrast but a different spatial frequency (Fig. 3E). As expected, Peak-C detected the optimal correction collar position, even under an XY-shuffled image, implying that the Peak-C algorithm ignored the biological structure, but the power spectrum of the XY-shuffled image shows a flat distribution (Fig. 3F). Contrary to our expectation that Peak-F reflects the fine structure as the spatial frequency, Peak-F also found the optimum position in the XY-shuffled images because the optimum correction collar provides the brightest images (Fig. 3G).

To demonstrate that the algorithm does not simply reflect the image’s brightness to determine the optimal correction collar position, we added saturated pixels to the benchmark image by increasing the pixel gain beyond the threshold without altering its spatial frequency (Fig. 4A). As the number of saturated pixels increased, the incorrect position was identified as the optimum using the Peak-C algorithm. In contrast, the Peak-F algorithm accurately found the optimum correction collar position even when 0.03% of pixels were saturated (Fig. 4B). These results along with those for the XY-shuffled cases (Figs. 3E-G) suggest that Peak-F does not simply look at brightness but also recognizes biological structures in the image.

### Peak-F reveals optical characteristics of the brain tissue from noisy images

The Peak-F algorithm is robust against noise and saturation (Figs. 1–4). Thus, we implemented Peak-F in a two-photon microscope system to test whether it was applicable to actual in vivo imaging. We observed samples 100, 200, 300, and 400 μm below the brain surface in both young (9-, 11-, 25-, and 25-weeks) and old (52-, 81-, 83-, and 93-weeks) mice (Fig. 5A). As expected, the deep area of old mice looked noisy, which may reflect the optical characteristics of the tissue such as RIs. We previously reported that our system can quantify RIs in any sample using the relationship between imaging depth and the correction collar angle[8]. Applying the Peak-F algorithm to detect the optimum correction collar angle in the noisy image, we calculated RIs in a deep area of aged mice. Figure 5B shows a relationship between the depth and optimum correction collar angle. Converting this relationship to RIs, we found a significant difference between young and old mice between 300–400 μm (Fig. 5C, 1.36 ± 3.88E-3 vs. 1.46 ± 5.99E-2, p = 0.04).

## Discussion

The ability of the proposed algorithm, Peak-F, to find the optimum correction collar position was compared to that of Peak-C. Specifically, we verified Peak-F’s performance on noise-added images (Fig. 2), binning images (Fig. 3), XY-random shuffled images (Fig. 3), and saturation-added images (Fig. 4). Then Peak-F was applied to two-photon microscopy to observe the brain of a mouse (Fig. 5).

Previously, we used Peak-C to determine the optimal correction collar position from images[8]. However, it was not applicable to noisy images. In contrast, Peak-F realized highly accurate calculations even in high-noise images by the Fourier transform and extracted 1–40 Hz. Since this study used 1024 pixels square for about 500 micrometers, 1–40 Hz corresponds to approximately 12.5 −500 mm. We believe that 40 Hz is sufficient to cover the soma size because Figs. 3 and 4 demonstrate that Peak-F reflects the fine biological structure.

The Peak-C algorithm calculates the optimum correction collar position according to the image’s contrast. Indeed, image processing such as noise addition, binning, and XY-random shuffling, does not affect the calculation. This algorithm compensates for the effect of spatial frequency in the contrast calculation by subtracting from multiple distant pixels (N = 2, 3, 5, 10, and 20). Because the impact of the spatial frequency on the analysis accuracy remained unclear, we also analyzed the contribution of each N (Fig. 3, Supplementary Fig. 1-A). The benchmark image gave homogeneous contributions for all frequency components at the optimal correction collar position, indicating that Peak-C tends to calculate the signals and high-frequency components such as noise equally. Hence, Peak-C is vulnerable to noise. In contrast, the contribution of the low-frequency components may depend on the object’s size. For example, the signal in the benchmark image contained sparse dendrites with about 1.5-μm diameter. In fact, in the artificially generated image with 1.5-μm diameter sparse bright spots, the contribution of the low-frequency component became saturated at ~20%, whereas the case of 10-μm diameter bright spots looked like as soma. That is, the 10-μm diameter bright spots had a much larger contribution of the low-frequency component while that of the high-frequency component was negligible (Supplementary Fig. 1B). Therefore, using only the low-frequency component for contrast calculations is sufficient for large structures such as cell bodies. Because actual living organisms have a wide variety of large structures, the contrast in every frequency band must be calculated to determine the optimal contrast value. This is an unrealistic high-cost computation. Alternatively, Peak-F is flexible and can handle the appearance of any structure, including noise, while maintaining a low computation cost.

The Peak-F algorithm uses the image frequency determined by the structure in the image for the calculation. Although it was hypothesized that it may not be suitable to determine the optimal image for XY-random shuffling, in which the shape of the power spectrum varies greatly, image brightness without a biological structure should contribute to the algorithm’s robustness. Peak-F chose the optimal image, which is probably because the image luminance determines the magnitude of power when a structure is not present. The XY Random Shuffle described earlier showed a structure change while maintaining brightness. We also compared the performance of each algorithm with saturation, in which the brightness changes while maintaining the structure. The Peak-C algorithm showed a weak performance, whereas the Peak-F algorithm was robust. Consequently, Peak-F improves the ability to make optimal image decisions for noise and saturation at a faster computational speed than Peak-C.

The amount of information in neighboring pixels is large because digital cameras use reduction optics[12]. Therefore, the auto-focusing technique used in a digital camera employs simple subtraction between adjacent pixels, which is sufficient to calculate the contrast of images[9]. This procedure is known as Brenner’s equation (Eq. 1). In contrast, microscopy employs magnifying optics, making microscopy susceptible to high-frequency components because the difference between neighbors is small and Brenner’s equation cannot adequately calculate the contrast. Therefore, we invented Peak-C. Unfortunately, Peak-C was adversely impacted by noise. Since the biological structures are concentrated in the low-frequency band, and excluding high-frequency components easily separates noise and frequency bands, the Fourier transform analysis in Peak-F is suitable to determine the optimal correction collar position.

The techniques used in adaptive optics can also correct spherical aberrations[13–15,2,16–20]. However, the optical path fluctuations must be calculated in advance since the brain tissue’s RIs are unknown. For example, embedded fluorescent beads with a known diameter and luminance at the given depth of the brain tissue are necessary to calculate the optical path fluctuations. Since a biological brain’s RIs may not be constant and vary with time and individual differences, accurate compensation requires implanting fluorescent beads in every trial, which is a time-consuming and invasive surgical procedure with many steps to obtain a single image. In contrast, our system is less invasive, and neither a high calculation cost nor additional optical devices are necessary. Additionally, there are no compulsory assumptions about the tissue’s RI.

Finally, we implemented the Peak-F algorithm for two-photon imaging of living mouse brains. The brains of old mice showed higher RIs than those of young mice at 300–400 μm. However, the Peak-C algorithm, which is sensitive to noise, had difficulty determining the optimal correction collar as old brain tissue tends to show more scattering in deeper areas. Hence, the noise-robust Peak-F algorithm may be useful not only for high-resolution observations of biological tissues but also for studying the physical properties of the brain. Since the spherical aberrant is the mismatching of RIs between water and tissue, the Peak-F algorithm can estimate timeseries changes in RIs. In the future, our system may be applicable to indirectly visualize the water dynamics in living biological tissues because the composition of water and lipid in biological tissues determines RIs.

## Acknowledgement

This work was supported by Ochanomizu University, RIKEN Center for Brain Science Institute, KAKENHI grants (18K14859, 20K15895), JST FOREST Program, Grant Number JPMJFR204G, Research Foundation for Opto-Science and Technology, Kao Research Council for the Study of Healthcare Science, The Japan Association for Chemical Innovation, and TERUMO LIFE SCIENCE FOUNDATION. We thank Hiroshi Hama, Reiko Takahashi, and Kana Namiki for providing the transgenic mice (YFP-H); Takako Kira, Koichi Imamura and all members in BOCC for assistance.

---

**Supplementary Figure 1.**
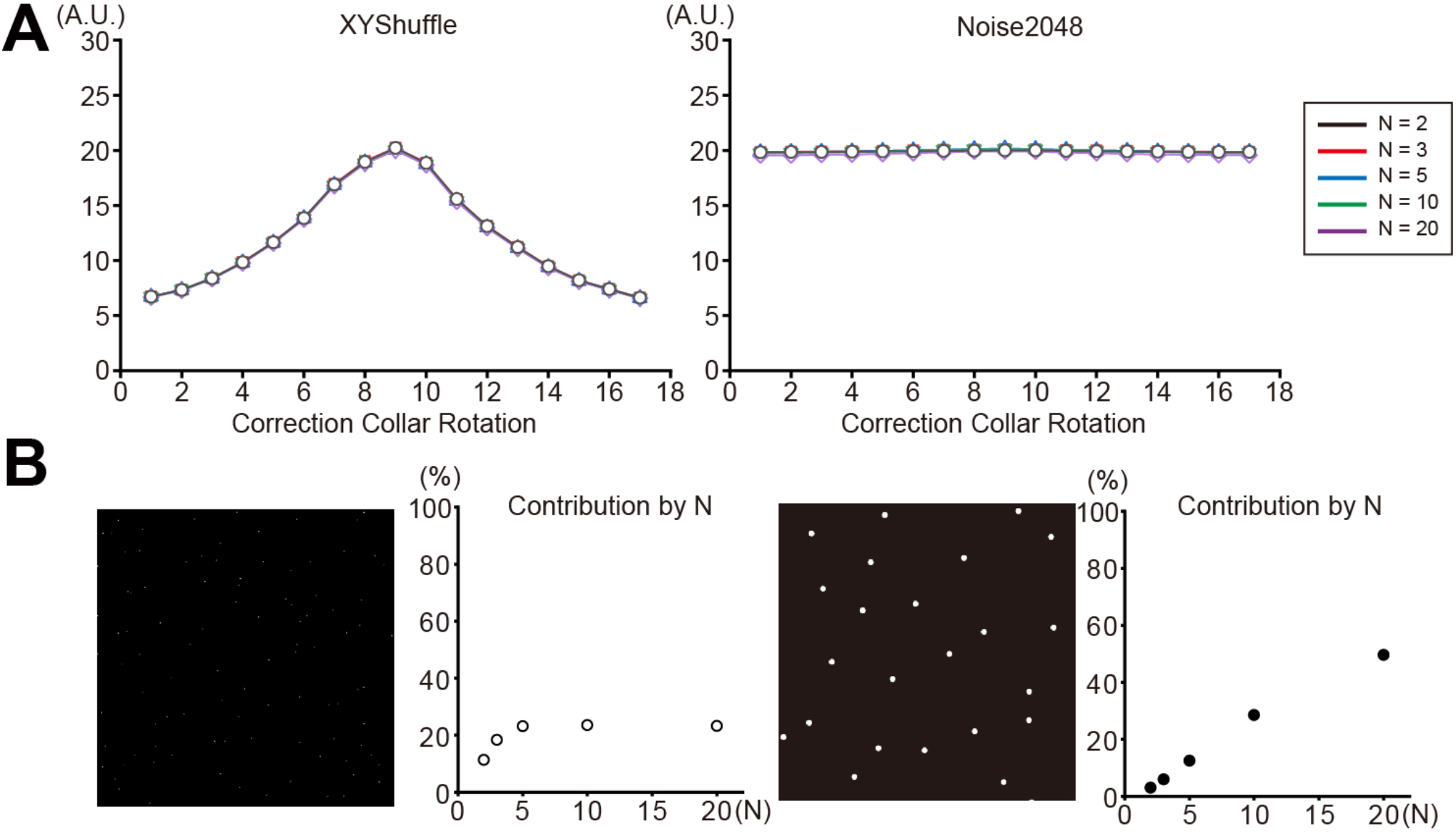
The effect of spatial frequency in the contrast calculation by subtracting from multiple distant pixels N A. The proportion of the contribution with each N (N = 2, 3, 5, 10, and 20) for the detection of the optimum correction collar position in XY-random shuffled images (left) and noise-added images (right). B. The proportion of the contribution with each N (N = 2, 3, 5, 10, and 20) at the optimum correction collar position in the artificially generated image with 1.5-μm diameter sparse bright spots (left) and 10-μm diameter bright spots (right).

